# Dynamic community detection reveals transient reorganization of functional brain networks across a female menstrual cycle

**DOI:** 10.1101/2020.06.29.178152

**Authors:** Joshua M. Mueller, Laura Pritschet, Tyler Santander, Caitlin M. Taylor, Scott T. Grafton, Emily Goard Jacobs, Jean M. Carlson

**Affiliations:** Interdepartmental Graduate Program in Dynamical Neuroscience, University of California, Santa Barbara, Santa Barbara, CA, USA; Department of Physics, University of California, Santa Barbara, Santa Barbara, CA, USA; Department of Psychological and Brain Sciences, University of California, Santa Barbara, Santa Barbara, CA, USA; Neuroscience Research Institute, University of California, Santa Barbara, Santa Barbara, CA, USA

**Keywords:** sex hormones, dynamic community detection, dense sampling, network flexibility

## Abstract

Sex steroid hormones have been shown to alter regional brain activity, but the extent to which they modulate connectivity within and between large-scale functional brain networks over time has yet to be characterized. Here, we applied dynamic community detection techniques to data from a highly sampled female with 30 consecutive days of brain imaging and venipuncture measurements to characterize changes in resting-state community structure across the menstrual cycle. Four stable functional communities were identified consisting of nodes from visual, default mode, frontal control, and somatomotor networks. Limbic, subcortical, and attention networks exhibited higher than expected levels of nodal flexibility, a hallmark of between-network integration and transient functional reorganization. The most striking reorganization occurred in a default mode subnetwork localized to regions of the prefrontal cortex, coincident with peaks in serum levels of estradiol, luteinizing hormone, and follicle stimulating hormone. Nodes from these regions exhibited strong intra-network increases in functional connectivity, leading to a split in the stable default mode core community and the transient formation of a new functional community. Probing the spatiotemporal basis of human brain–hormone interactions with dynamic community detection suggests that ovulation results in a temporary, localized patterns of brain network reorganization.

**Author Summary:** Sex steroid hormones influence the central nervous system across multiple spatiotemporal scales. Estrogen and progesterone concentrations rise and fall throughout the menstrual cycle, but it remains poorly understood how day-to-day fluctuations in hormones shape human brain dynamics. Here, we assessed the structure and stability of resting-state brain network activity in concordance with serum hormone levels from a female who underwent fMRI and venipuncture for 30 consecutive days. Our results reveal that while network structure is largely stable over the menstrual cycle, there is temporary reorganization of several largescale functional brain networks during the ovulatory window. In particular, a default mode subnetwork exhibits increased connectivity with itself and with regions from temporoparietal and limbic networks, providing novel perspective into brain-hormone interactions.

## Introduction

The application of network science techniques to the study of the human brain has revealed a set of large-scale functional brain networks that meaningfully reorganize both intrinsically and in response to external task demands [1]. One technique, dynamic community detection (DCD), has emerged as a powerful tool for conceptualizing and quantifying changes in mesoscale brain network connectivity patterns by identifying sets of nodes (communities) with strong intra-community connections [2] to enable identification of communities that persist or change over time. DCD complements other statistical approaches used in fMRI data analysis by identifying when functionally coupled brain regions undergo sufficiently large changes in connectivity to warrant re-assignment to separate functional communities. Additionally, this method provides an interpretable summary of whether strongly connected sets of brain regions undergo transient, but significant, changes that could be missed when time-averaging data within and between sessions.

This method is particularly suited for examining relationships between brain dynamics and physiological variables that vary over relatively short time scales, such as sex hormone fluctuations over the human menstrual cycle. A typical cycle, occurring every 25–30 days, is characterized by significant rises in estradiol (~12-fold) and progesterone (∼800-fold), both of which are powerful neuromodulators that have a widespread influence on the central nervous system [3]. Converging evidence from animal studies has established sex hormones’ influence on regions supporting higher-order cognition, including the prefrontal cortex (PFC) and hippocampus [4, 5]. Within these regions, fluctuations in estradiol enhance spinogenesis and synaptic plasticity while progesterone largely abolishes this effect [6, 7]. Importantly, sex hormones are expressed broadly throughout the cerebellum and cerebrum, suggesting that whole-brain effects might be observed beyond the regions targeted in these studies.

Human neuroimaging studies have demonstrated that sex hormones influence brain activity across broad regions of cortex [8, 9]. Additionally, a handful of studies have demonstrated that menstrual cycle stage uniquely alters resting-state functional connectivity (rs-fc) [10–13]. However, these studies typically involve group-based or sparse-sampling (2–4 time points) designs that are unable to capture transient day-to-day relationships between sex hormones and functional brain dynamics, and this relatively low temporal resolution has led to inconsistencies in the literature [14]. Therefore, new approaches are needed that can address these spatial and temporal limitations, as doing so will provide novel perspectives on human brain–hormone interactions.

Recently, Pritschet et al. applied a “dense sampling” approach [15, 16] to a naturally-cycling female who underwent 30 consecutive days of brain imaging and venipuncture to capture rs-fc variability over a complete menstrual cycle (Fig 1). The authors found day-to-day fluctuations in estradiol to be associated with widespread increases in rs-fc across the whole brain, with progesterone showing an opposite, negative relationship. Using time series modeling and graph theoretical analysis, they also found that estradiol drives variation in topological network states, specifically within-network connectivity (global efficiency) of default mode and dorsal attention networks that encompass regions rich with estrogen receptors (ER). These findings have important implications for the field of network neuroscience where dense-sampling, deep-phenotyping approaches have emerged to aid in understanding sources of intra/inter-individual variability in functional brain networks over days, weeks, months, and years [16–18].

**Fig 1.**
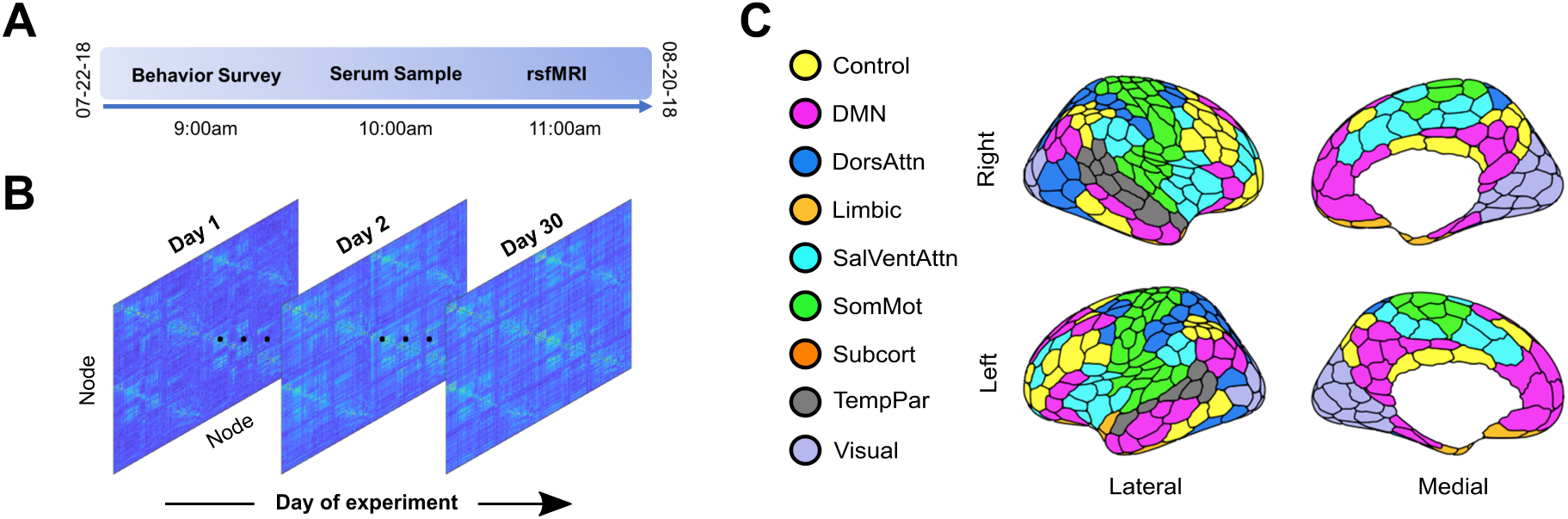
28andMe dataset. **A**. Subject LP (naturally cycling female, age 23) participated in a month-long “dense sampling” experimental protocol to provide a multimodal, longitudinal dataset referred to as 28andMe [19]. The subject completed daily assessments of diet, mood, and sleep, provided blood for assessment of serum hormone concentrations, and underwent a 10 minute resting-state fMRI scan. **B**. For each resting-state scan, functional connectivity matrices were constructed by calculating the pairwise mean magnitude-squared coherence between each region from the entire 10-minute scan. The result is a 415 × 415 × 30 data structure, in which each entry indicates the coherence between two nodes on a given day. **C**. The brain was parcellated into 415 regions and regions were assigned to one of nine networks based on previously identified anatomical and functional associations [20]. Colors indicate regional network membership.

Pritschet and colleagues’ approach identified node-averaged trends in rs-fc changes within canonical functional networks across the cycle, but questions remain regarding whether and where functional reorganization takes place between large-scale networks. As changes in edge weight can result in the formation of functional “communities” not captured by traditional rs-fc methods, complementary approaches are needed to characterize trends in brain connectivity at intermediate spatial and temporal scales. Examining mesoscale networks has further revealed fundamental principles of functional brain networks, such as the modular, integrated architecture underpinning flexible task performance [21, 22]. Additionally, a better understanding of mesoscale connectivity may provide an avenue for improving personalized medicine by increasing the efficacy of targeted therapeutic interventions [23].

Here, we applied DCD to examine whole-brain dynamics in relation to sex hormone fluctuations across a menstrual cycle. Our results reveal that a stable set of “core” communities persist over the course of a menstrual cycle, primarily consisting of nodes belonging to distinct *a priori* defined functional–anatomical networks, namely visual, somatomotor, attention, default mode, and control networks. Though these core communities were largely stable, nodes from limbic, subcortical, attention, and control networks changed community affiliation (referred to as flexibility) at higher rates than expected compared to a null hypothesis. DCD also identified a transient split of the DMN core into two smaller subcommunities concurrent with peaks in estradiol, luteinizing hormone (LH), and follicle stimulating hormone (FSH) levels defining the ovulatory window. This community split was driven by strong increases of within-network integration between prefrontal nodes of the DMN, which subsided immediately after the ovulatory window. The default mode, temporoparietal, limbic, and subcortical networks also exhibited significantly increased flexibility during ovulation, suggesting a role for estradiol, LH, and FSH in regulating localized, temporary changes in regional connectivity patterns. Taken together, while a large degree of functional brain network stability was observed across the menstrual cycle, peaks in sex hormones over the ovulatory window resulted in temporary brain network reorganization, suggesting sex hormones may have the ability to rapidly modulate rs-fc on shorter time scales than previously documented.

## Results

A single female underwent brain imaging and venipuncture for 30 consecutive days. For each session, the brain was parcellated into 400 cortical regions from the Schaefer atlas and 15 subcortical regions from the Harvard-Oxford atlas (Fig. 1C) and 415 × 415 functional association matrices were constructed via magnitude-squared coherence [20]. Dynamic community detection was applied to this data, revealing a stable set of communities that persist over the course of a menstrual cycle. However, significant transient changes in community structure occurred within the default mode network during the ovulatory window concomitant with peaks in estradiol, luteinizing hormone, and follicle stimulating hormone.

### Stable functional cores persisted over the course of one menstrual cycle

The degree to which functional brain network connectivity changes over the course of a human menstrual cycle has yet to be fully characterized. Here, dynamic community detection (also referred to as multislice or multilayer modularity maximization [24]) consistently identified four functional communities that were largely stable in a naturally cycling female over 30 consecutive days. In this context, “community” refers to a set of nodes whose intra-set connections are significantly stronger than would be expected when compared to an appropriate null model. A representative example of this consensus temporal community structure (the community designation that best matches the output of 50 runs of the non-deterministic community detection algorithm) is shown in Fig. 2C. This structure was conserved over a range of community detection parameter values that, roughly speaking, must be defined to set the “spatial” and “temporal” resolutions of community identification (see Methods for detailed description). Across all temporal resolutions considered here, consensus community partitions with a spatial resolution parameter 0.975 *≤ γ ≤* 1.01 possessed exactly four communities.

**Fig 2.**
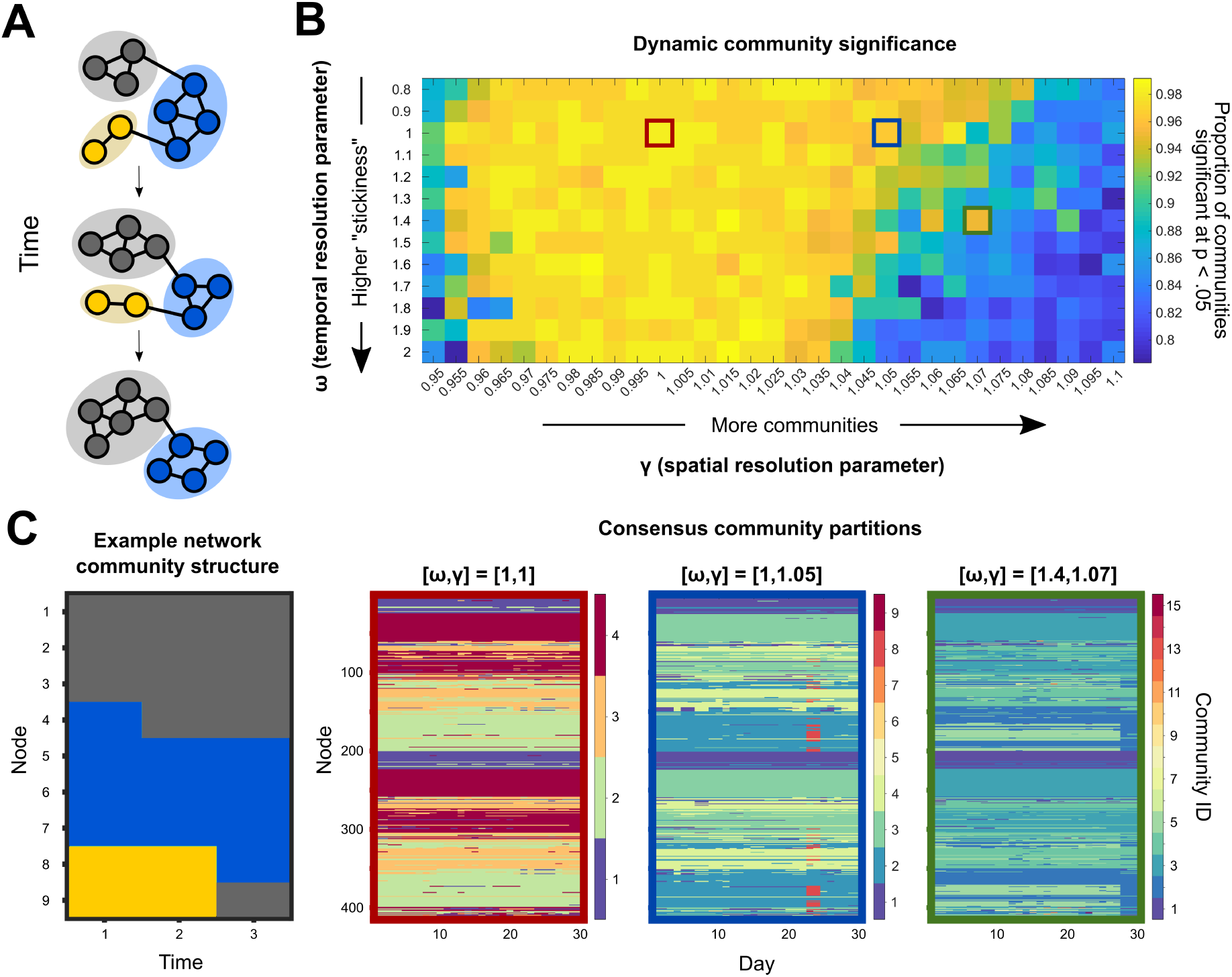
Dynamic community detection identified changing modular structure over time at multiple scales. **A**. A toy network example illustrates the dynamic community detection algorithm. For each time point, every node is assigned to a community so as to maximize the strength of intra-community connections relative to inter-community links while also taking community assignments over time into account (Eq. 1). In this case, three communities are identified and denoted by color. **B**. To assess temporal structure in the 28andMe resting-state fMRI data, community assignments were calculated for a range of parameter values. In this procedure, two parameters, *ω* and *γ*, specify the temporal and spatial scales of analysis, respectively. After performing 50 runs of the community detection algorithm for each parameter combination, the statistical significance of each community partition relative to a random null model was calculated. The color for each entry in the heat map indicates the proportion of communities at that parameter combination which are significant at the *p <* .05 level. **C**. Consensus partition structure varied according to the choice of resolution parameters. The example network community structure (left) changes at each time point, with node community assignment given by color on the y-axis and time indicated on the x-axis. For three different parameter combinations (outlined in red, blue, and green, respectively), the consensus partitions varied in the total number of communities identified, ranging from four to fifteen, with more communities identified when the temporal resolution was low and the spatial resolution was high.

For the standard parameter choice (temporal and spatial resolution parameters both set to 1), the four identified communities had distinct compositional characteristics. These communities were largely bilaterally symmetric, with analogous brain regions in each hemisphere assigned to the same community 71% of the time. The four communities correspond roughly to a visual core, a somatomotor-attention core, a default mode core, and a control core. The compositions of these four communities are shown in Fig. 3A. The composition value was calculated by summing the total number of instances in which a node belonging to an *a priori* functional-anatomical network [20] also belonged to the community identified in the consensus community partition.

**Fig 3.**
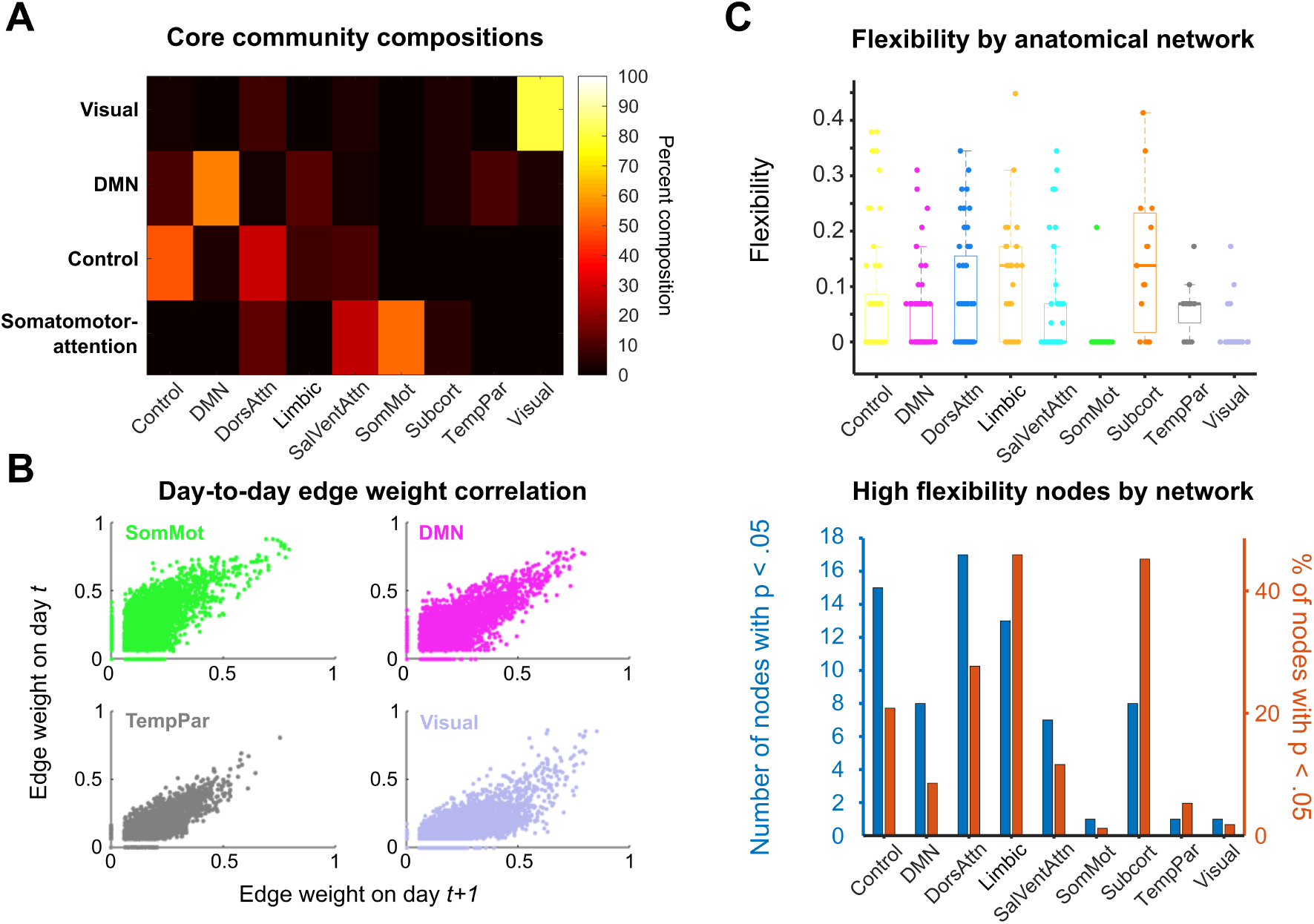
Dynamic community detection uncovered stable cores across a complete menstrual cycle. **A**. Four core communities (y-axis) were consistently identified in the 28andMe dataset across spatial and temporal resolution parameter values. For these parameter combinations, the compositions of the visual, default mode, control, and somatomotor-attention network cores are shown as a heat map, with color corresponding to the percentage of nodes in a community belonging to a functional-anatomical network. **B**. The four networks that constituted the hubs of the core communities possessed stable pairwise connectivity between nodes across days. Scatter plots show the day-to-day correspondence between edge weights for all of the nodes of the somatomotor, default mode, temporoparietal, and visual networks on days *t* and *t* + 1. These network edges had Pearson correlation coefficients of 0.379, 0.573, 0.590, and 0.538, respectively. **C**. The subcortical, limbic, and dorsal attention networks exhibited the highest median node flexibility. Top: Normalized flexibility values for each node over the entire cycle are plotted as points, with color indicating network affiliation. Thick horizontal lines on box plots indicate median values. A flexibility value of 1 indicates that a node changes community assignment at each possible time point, whereas a value of 0 indicates that the node never changes community assignment. Bottom: A 95% cutoff value is calculated using the flexibility values for each node over all 50 community detection runs. For each functional-anatomical network, the blue bar indicates the number of nodes belonging to that network which have flexibility values above the cutoff threshold. The red bars indicate the proportion of nodes in each network that surpass the cutoff value (i.e. the value for each blue bar is normalized by the number of nodes in the network). Once again, limbic, subcortical, dorsal attention, and control networks contained the highest proportion of highly flexible nodes.

The core communities identified here were named based on the highest representation of nodes belonging to *a priori* functional networks. The visual core was 81% composed of visual network nodes and had an average size of 51.5 nodes per day. The somato-attention core was composed of 52% somatomotor, 27% salience-ventral attention, and 13% dorsal attention network nodes and had an average size of 134.1 nodes per day. The default mode core consisted of 56% DMN nodes and approximately 10% of each control, limbic, and temporoparietal network nodes and contained 132.9 nodes on average per day. Finally, the control core consisted of 48% control and 28% dorsal attention network nodes and contained 96.5 nodes on average per day. Importantly, for all parameter combinations in which four communities were detected, the composition of these communities was consistent (Supplementary Information). These community partitions were also stable across the entire menstrual cycle. Specifically, 315 of the 415 nodes (75.9%) did not change community affiliation across the 30-day experiment.

Taken together, these results suggest the presence of a stable solution to the dynamic community detection algorithm and a reliable coarse-grained community architecture present in the data. In several functional–anatomical networks, there was little to no modification of network architecture over time; for instance, greater than 85% of nodes in each of the so-matomotor, default mode, temporoparietal, and visual networks did not change community affiliation over the entire menstrual cycle. The strong day-to-day correlations between edge weights in these networks (Fig. 3B) reinforce the existence of these stable cores.

### Functional-anatomical networks exhibiteddistinct patterns of flexibility

Though network community structure was stable over a complete menstrual cycle when classifying nodes into four communities, specific nodes did change community affiliation at levels above chance when modifying the sensitivity of the community detection algorithm. Specifically, when *γ*, the spatial resolution parameter, was increased, the dynamic community detection algorithm subdivided the four core communities into smaller communities, providing a finer-grained classification of subnetwork structure. At an intermediate parameter combination (*ω* = 1, *γ* = 1.05), nine communities significant at the p *<* .05 level were identified over the course of the experiment, as visualized in Fig. 2C (blue outlines). The subsequent analysis uses community partitions at this parameter combination, but the results were consistent across a range of neighboring parameter values (Supplementary Information).

This “higher-resolution” partition revealed trends in functional organization over time that were not observable with coarser partitions. First, inspecting the median flexibility value, or the proportion of times a node changed community affiliation out of the total possible number of changes, demonstrates that functional–anatomical networks possessed distinct flexibility distributions (Fig. 3C, top). The limbic, subcortical, dorsal attention, and control networks were overrepresented in terms of highly flexible nodes relative to a null hypothesis (Fig. 3C, bottom).

Fine-scale community reorganization occurred on experiment day 23 and persisted until day 25, as illustrated in Fig. 4A. Across these days, 62 nodes belonging to the default mode core community split from the default mode core community to transiently form a small, strongly connected community. This was the only large-scale reorganization event detected during the experiment, as indicated by the nodal flexibility values illustrated in Fig. 4B. Global flexibility was significantly higher (Wilcoxon rank-sum test, *p <* .05) during ovulation (days 23–25) than during follicular or luteal phases (days 11–22 and 26–10, respectively). Specifically, global mean flexibility during the ovulatory window was 0.142, whereas flexibility during follicular and luteal phases was 0.049 and 0.050, respectively.

**Fig 4.**
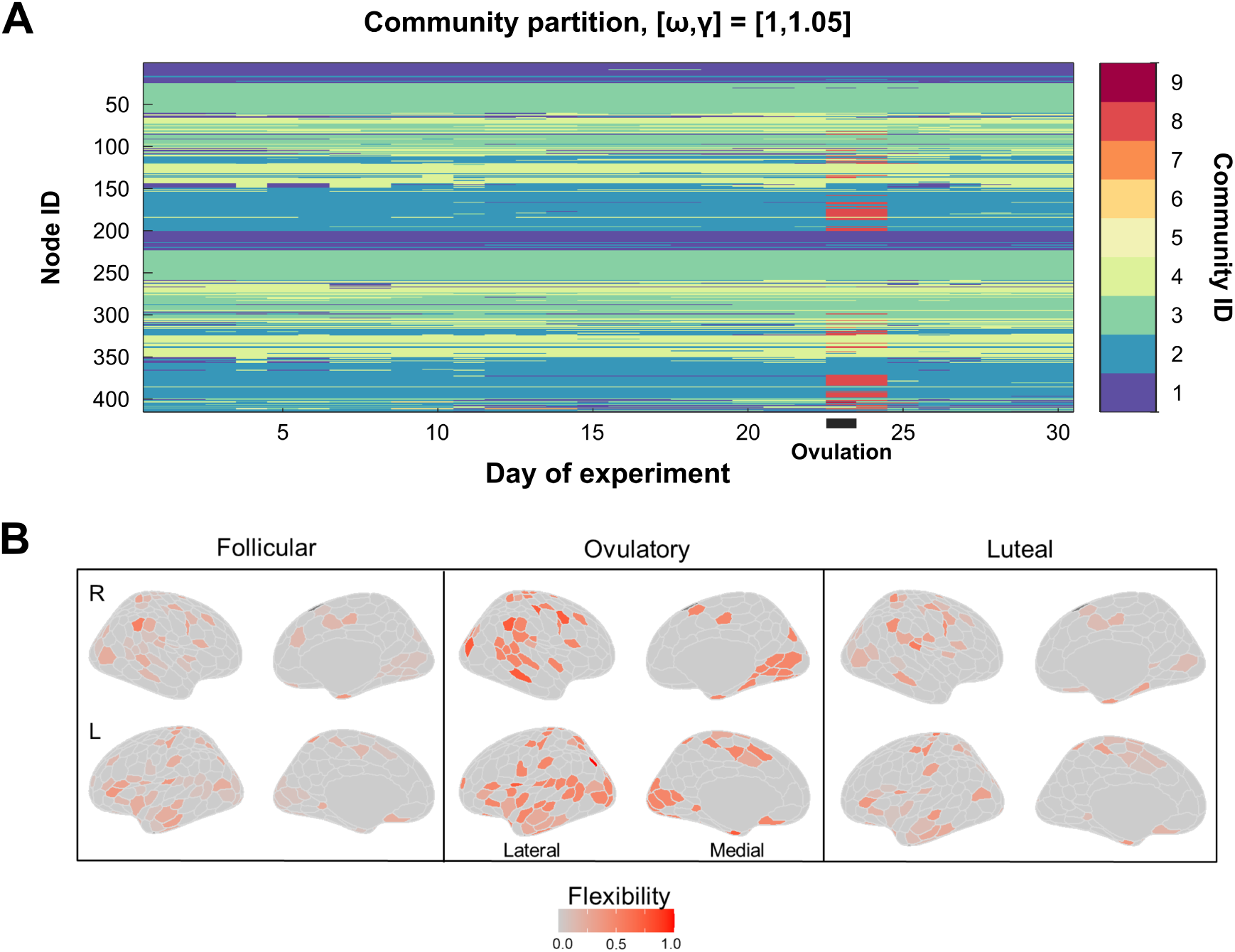
Fine-grain community partitioning revealed a bifurcation in the default mode core during ovulation. **A**. When the spatial resolution parameter (which alters the size of communities identified by dynamic community detection) was increased from the standard value, the four core communities identified previously were subdivided into smaller subcommunities (reproduced from Fig. 2C). Here, a split in the default mode core community (light blue) appeared at day 22 (red-orange), concomitant with ovulation and a spike in sex hormones. This community (red) rejoined the default mode core on day 25. For illustrative purposes, only the consensus partition for one parameter value is shown, but this trend was consistent across nearby parameter combinations (Supplementary Information). **B**. Shown are flexibility values for each node by menstrual cycle phase. Color in each region indicates flexibility value, with hotter colors indicating higher values. The following days of the experiment corresponded to the phases of the menstrual cycle: follicular, days 11-22; ovulatory, days 23-25; luteal, days 1-10 and 26-30. Flexibility values are noticeably higher in many regions from the temporoparietal, limbic, subcortical, and default mode networks during the ovulatory phase compared to the follicular and luteal phases.

Notably, 31 (50%) of the nodes in the community that emerged during the ovulatory window belonged to the DMN, 12 nodes (19%) belonged to the temporopari-etal network, and 7 (11%) were subcortical regions (as defined by functional–anatomical atlases [20, 25], Fig. 5A). The functional–anatomical network memberships of the node–node pairs exhibiting the strongest increases in coherence (top 5%) indicated that enhanced connectivity between DMN nodes drove this community split, as opposed to DMN nodes being “converted” to a new community via increased connectivity to non-DMN regions (Supplementary Information). More specifically, nodes within prefrontal regions belonging to DMN subnetwork B drove this reorganization event, as 118 of the 371 (32%) strongest increases in coherence occurred between nodes in this subnetwork.

**Fig 5.**
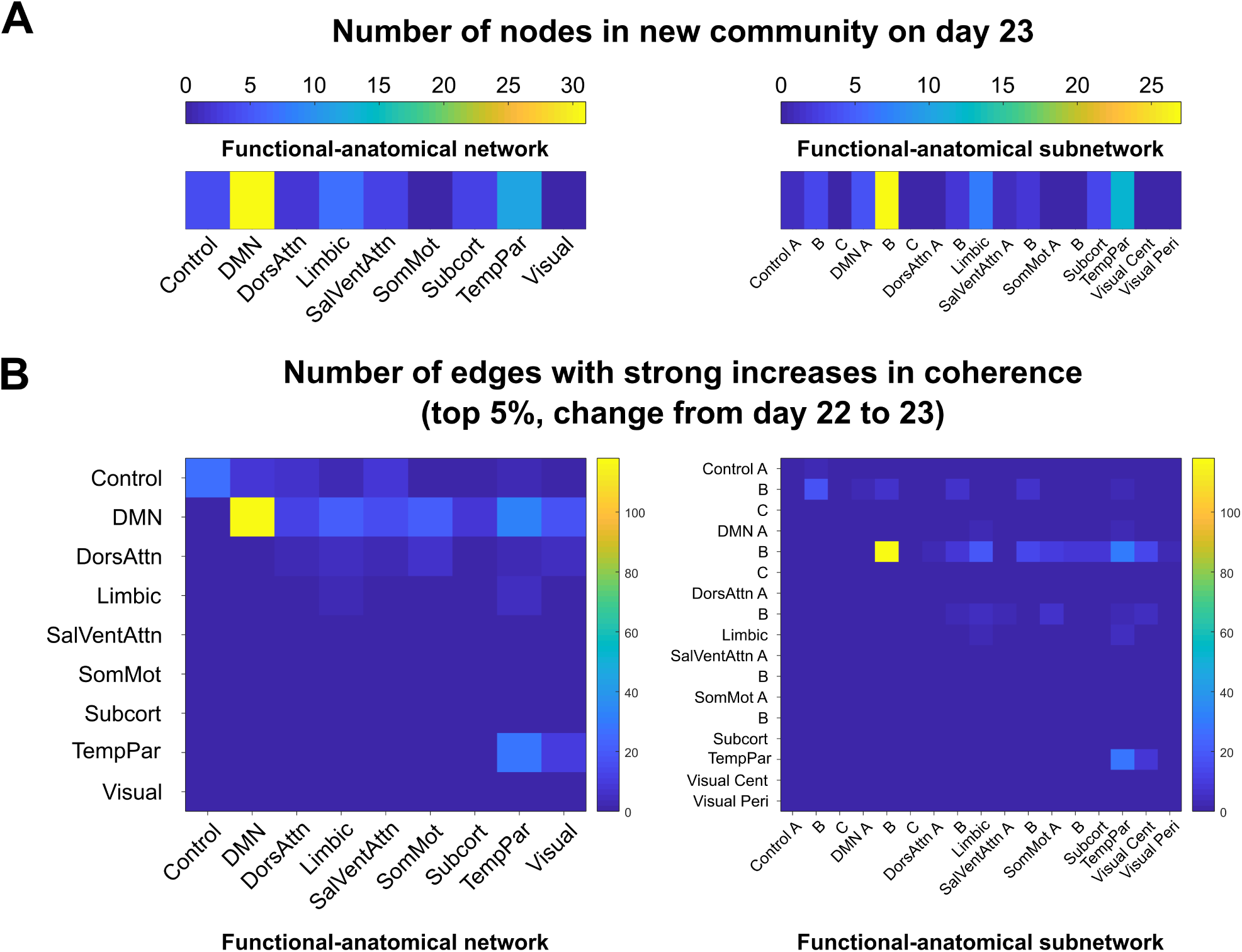
Nodes in a default mode subnetwork drove community bifurcation via strong increases in coherence. **A**. The newly formed functional community on day 23 and 24 contained 62 nodes that belonged to the community on both days. The functional-anatomical network and subnetwork affiliations of these nodes are shown on the left and right, respectively. The new community contained 31 DMN nodes (50%), 12 temporoparietal nodes (19%), and 7 subcortical nodes (11%). **B**. The edges that exhibited large weight changes from day 22 to day 23 (top 5% of changes, left) were predominantly within-network connections between DMN network nodes (118/371). Examining subnetwork structure reveals that all of the strongly enhanced connections between nodes in the DMN belonged to subnetwork B, indicating that this subnetwork, which consists of regions in prefrontal cortex, drove the default mode core community bifurcation at ovulation (Supplemental Information).

### Network reorganization timing coincided with peaks in hormone levels during ovulation

Mean flexibility of each network over a 5-day sliding window is depicted in Fig. 6A. The DMN, temporopari-etal, subcortical, and limbic networks exhibited peaks in flexibility at days 23 and 24 of the experiment, coin-cident with the peaks in estradiol, LH, and FSH which are a hallmark signals of the ovulatory window (Fig.6B). To determine whether the bifurcation of the default mode core community was significantly associated with sex hormones, we compared functional–anatomical network flexibility values to serum hormone levels.

**Fig 6.**
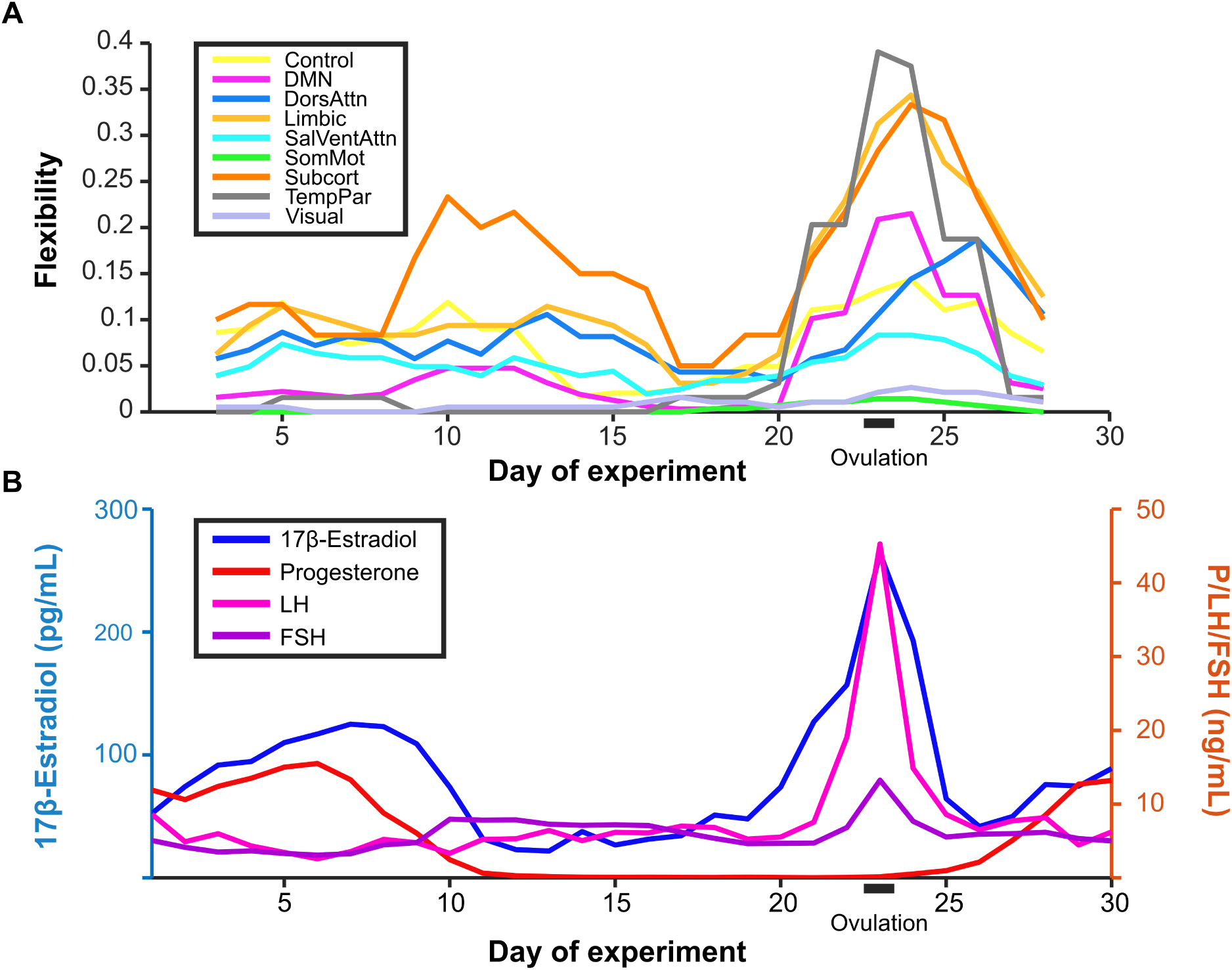
Community reorganization was temporally localized to ovulation. Changes in community assignment (**A**) were coordinated and closely tracked the timing of spikes in serum hormone concentrations (**B**). Prior to day 20 of the experiment, all networks except for the subcortical network exhibited low baseline rates of flexibility (mean = 0.04). However, several networks exhibited sharp increases in flexibility between days 20 and 26, indicating brain-wide functional reorganization during the ovulatory window. The pattern of flexibility shown here corresponds to the network reorganization observed for dynamic community detection performed with the parameter combination *ω* = 1, *γ* = 1.05 (blue outline in Fig. 2). To note, flexibility is calculated over a 5 day sliding window.

To assess the temporal relationship between network flexibility values and sex hormones, cross- covariance structure between each time series was calculated. The control, default mode, limbic, salience/ventral attention, subcortical, and temporoparietal networks had maximum cross-variance values greater than 0.6 (where maximum value of 1 indicates fully shared covariance structure and 0 indicates no covariance) with estra-diol, which were significant when compared to cross-covariance values for a null model of time- permuted estradiol levels (Bonferroni-corrected at p *<* .05). Each network except for the control and attention networks had maximum cross-variance values greater than 0.6 with LH as well (permutation test, p *<* .05 after Bon-ferroni correction). In each case, maximum cross-covariance values occurred at lags less than 2 days and no other significant cross-covariance structure existed, indicating that most functional communities exhibited changes in composition concurrent with significant rises in estradiol and LH levels.

## Discussion

In this study, we applied DCD to data from a densely sampled female who underwent 30 consecutive days of brain imaging and venipuncture to investigate the extent of intrinsic spatiotemporal functional reorganization over a menstrual cycle. We identified four stable community cores across the cycle, represented here as visual, somatomotor, default mode, and control network cores; interestingly, the exception to this stability occurred simultaneously with peaks in estradiol, LH and FSH. During this event, we observed a transient reorganization of the default mode core into a newly formed community, as well as increases in nodal flexibility among prefrontal, limbic, and subcortical nodes. Taken together, our results suggest that the interplay between the nervous and endocrine systems over a menstrual cycle result in temporary, localized patterns of brain network reorganization occurring during ovulation. These results highlight DCD as a new avenue for investigating the intricate relationship between sex hormones and human brain dynamics.

### Dynamic community detection characterizes network-specific functional stability across a menstrual cycle

Dense-sampling, deep-phenotyping studies offer new ways to investigate intra/inter-individual variability in functional brain networks by identifying features of rs-fc that are stable traits within an individual or change in conjunction with biological factors and state-dependent variables [16, 18]. Recent dense-sampling studies have shown that frontoparietal regions/networks exhibit high degrees of intra-individual rs-fc stability while also being characteristically unique across individuals, suggesting that these higher-order regions may be especially critical for uncovering individual differences in brain function and improving applications into personalized medicine [18, 26]. Our findings provide new insight towards the ongoing explorations into stability within functional brain networks. In this dataset, frontal control and DMN nodes exhibited high day-to-day connection weight correlations and low propensity to change functional community membership over the experiment (Fig. 3), while, on average, somatomotor, temporoparietal, visual, and salience/ventral attention networks were also largely stable. Therefore, our results align with previous research suggesting both a high degree of network stability in resting-state networks over a menstrual cycle [14] and in individuals over time [16, 18, 26].

In conjunction with this observed stability, network-specific changes in functional community organization were also identified. Control subnetwork C, encompassing posterior cingulate cortex/precuneus regions, was the most flexible functional subnetwork identified, with 10 of the 12 nodes exhibiting significantly higher than expected flexibility (i.e. how often a node switches community affiliation, see Supplementary Information). Limbic and subcortical networks displayed intermediate levels of flexibility. Regions from these systems receive input from and project to many cortical areas and are implicated in functions such as sensorimotor integration via the cortico-basal ganglia-thalamo-cortical loop [27]; therefore, the high degree of flexibility observed here may reflect the tendency of these systems to serve as relays between functionally segregated communities.

Particular changes in rs-fc were significantly related to the sharp rises in sex hormones seen across the ovulatory window. During this time, we observed a spatially-specific transient reorganization of the DMN, during which nodes from the temporoparietal, limbic, subcortical, and default mode networks split from the default mode core to form a short-lived community (2 days) before rejoining the original core community. Using time-lagged analyses, Pritschet and colleagues previously reported that within-network connectivity of the DMN was regulated by previous states of estradiol [19]. Here, we expand on this finding and identify a subnetwork of the DMN that is likely driving this reorganization. Notably, regions constituting this new community are located in PFC, an area exquisitely sen-sitive to sex steroid hormones [28] where, for instance, nearly 50% of pyramidal neurons in the dorsolateral PFC (dlPFC) express ER-alpha [4]. Together, this presents the possibility that endocrine signaling may, in part, regulate intrinsic brain dynamics within the frontal cortex.

### Neurobiological interpretations of sex hormones on PFC function

Cross-species investigations have established estrogen’s ability to shape the PFC [9,28–31]. In rodents, estradiol increases fast-spiking interneuron excitability in deep cortical layers [32]; in non-human primates, estradiol treatment increases dendritic spine density in dlPFC neurons [33] and this potentiation is observed only if the treatment is administered in the typical cyclical pattern observed across a menstrual cycle. In parallel, human brain imaging studies have implicated estradiol in enhancing the efficiency of PFC-based circuits. In cycling women performing a working memory task, PFC activity is exaggerated under low estradiol conditions and reduced under high estradiol conditions [9]. Similarly, when estradiol declines across the menopausal transition, working-memory related PFC activity becomes more exaggerated despite no differences in task performance [31]. Examining rs-fc across the cycle, Petersen and colleagues found that women in the late follicular stage (near ovulation) showed increased coherence within the default mode and executive control networks compared to those in luteal stages [10]. Our findings extend this body of work by demonstrating that dlPFC nodal flexibility tracks significantly with sharp increases in estradiol and LH across the cycle, which may support the brain’s ability to reorganize at the mesoscale level.

This tight temporal coupling highlights the potential for a mechanistic link between endocrine signaling and large-scale network reorganization. While future multimodal brain imaging studies are needed to establish this link, one possible neurobiological mechanism of action may be through estradiol’s interaction with the dopaminergic system. For instance, the PFC is innervated by midbrain dopaminergic neurons that enhance the signal-to-noise ratio of PFC pyramidal neurons and drives cortical efficiency [34]. In turn, estradiol enhances dopamine release and modifies the basal firing rate of dopaminergic neurons, providing one explanation for how alterations in estradiol could impact cortical efficiency.

### Implications for cognition and disease

Several studies have begun utilizing DCD to relate “task-free” and “task-based” functional network reorganization to cognitive performance. High levels of nodal flexibility have been associated with enhanced performance on working memory tasks [35], improved learning of a motor task [36], and visual cue learning [37]. In each study, flexibility was associated with performance in regions known to underlie each task, implicating frontal, motor, and visual cortical cortices and subcortical structures such as thalamus and striatum. Notably, similar associations were not observable when analyzing these experiments through region-based activation patterns alone, indicating that temporal organization of brain-wide functional activity (e.g. dynamic community structure) may provide important information related to cognitive functioning that might be missed with traditional analyses.

Indeed, Mattar et al. used DCD to characterize cognitive systems like those defined here [20] in a 64-task battery, demonstrating that functional networks fluidly reconfigure to form new cohesive communities under different task settings [38]. Similar work has revealed that primary motor, visual, and auditory regions typically participate in a single or a small number of functional networks during various tasks, whereas “hub” regions in frontal cortex, including precuneus and posterior cingulate gyrus participate in multiple functional networks [39]. Together, these studies indicate that network-specific temporal reconfiguration of functional connectivity has implications for a wide variety of cognitive functions. While whole-brain activity patterns during task-free states differ from that of goal-directed cognitive states, the capacity for the brain to fluctuate between integrated and segregated (modular) states at rest allows for rapid and efficient transitions to various task states [40–42]. Here, we leverage these techniques to characterize the brain’s response to both subtle and pronounced hormonal changes typical of a menstrual cycle.

Highly flexible nodes were identified in precuneus and posterior cingulate gyrus, with changes in community affiliation occurring simultaneously with sharp peaks in estradiol and LH levels, raising the possibility that hormonal fluctuations could also be associated with task-based network reorganization. For instance, if high levels of estradiol increase nodal flexibility among hub regions in the PFC, one would predict that performance on PFC-dependent tasks will improve. Further, pregnancy—a period of profound hormonal change —leads to long-lasting gray matter reductions in regions within the default mode network [43]. Therefore, future work examining whether task-based functional brain networks undergo transient changes in flexibility and community structure both across the menstrual cycle and during other hormonal transition periods, and whether this impacts cognitive performance, will be imperative.

Examining how large-scale brain networks are disrupted between healthy and patient populations may enhance our understanding of neurological conditions [44]. Notable intrinsic connectivity differences within the DMN are observed among individuals with depression [45] and Alzheimer’s disease [46]– two conditions that display a sex-skewed prevalence towards women [47]. Recent studies have applied DCD methods to characterize functional brain network reconfigurations in different disease states: region-specific flexibility at rest has been linked to symptom severity in autism spectrum disorder [48] and a recent investigation used DCD to associate pronounced community reorganization during seizures with poorer surgical outcomes [49]. Here, using similar methods, we demonstrate that high estradiol days are associated with significant reorganization of the default mode network and increased flexibility of several brain networks. Understanding the relationship between brain network reconfiguration (time-varying communities) and the endocrine system (dynamic fluctuations in sex hormones) may offer new ways to understand complex neurological conditions, especially those with pronounced sex differences in disease prevalence.

### Limitations and future directions

The following limitations should be taken into consideration. First, this study involved densely sampling a single female over one complete menstrual cycle, hindering our ability to generalize these findings to other individuals. Therefore, it is critical for this approach to be extended to a larger and more diverse set of women to establish the consistency of these results while taking individual differences into consideration. Second, we used a well-established group-based atlas to mitigate the limitations inherent to a single-subject design and improve generalizability [20]. However, recent work has demonstrated that group-based atlases can lead to loss in individual-level specificity and overlook meaningful spatial reconfigurations in parcellations themselves [50]. Future work using an individual-derived atlas is needed to confirm whether these results are stable across various parcellation applications. Finally, an ongoing debate in network neuroscience surrounds testretest reliability and what constitutes a “substantial” amount of data per individual. While some studies suggest that large amounts of data (*>* 20 minutes) is needed [18], others contend that shorter durations (5–15 minutes) of sampling is sufficient to achieve reliability [17, 51]. Repeating this experiment under longer scanning durations (*>*10 minutes per day) will be critical for exploring the degree of network stability across the menstrual cycle.

## Conclusion

In sum, we demonstrate that resting-state functional connectivity is largely stable within an individual over the course of a complete menstrual cycle. The exception to this stability occurs around the ovulatory window, during which peaks in sex hormones result in temporary patterns of brain network reorganization largely localized within areas of the default mode network. Historically, brain-level phenomena resulting from hormone fluctuations have been treated as an unwanted source of variance in population studies and, consequently, studies of this relationship are sparse and underpowered. This work demonstrates that dynamic network methods can reveal important, transient effects of sex hormones that may be overlooked by traditional approaches and provides a novel template for examining the nature of human brain–endocrine relationships.

## Methods

### 28andMe experimental protocol

Data was collected and preprocessed as reported in [19]; methods briefly reproduced here. The participant was a right-handed Caucasian female, aged 23 years for duration of the study. The participant had no history of neuropsychiatric diagnosis, endocrine disorders, or prior head trauma. She had a history of regular menstrual cycles (no missed periods, cycle occurring every 26–28 days) and had not taken hormone-based medication in the 12 months prior to the study. The participant gave written informed consent and the study was approved by the University of California, Santa Barbara Human Subjects Committee.

The participant underwent daily testing for 30 consecutive days, with the first test session determined independently of cycle stage for maximal blindness to hormone status. The participant began each test session with a daily questionnaire (9:00am) followed by a *±* time-locked blood sample collection 10:00am (±30 min). Endocrine samples were collected, at minimum, after two hours of no food or drink consumption (excluding water). This was followed by a one-hour MRI session (11:00am) consisting of structural and functional MRI sequences. To note, the participant refrained from consuming caffeinated beverages before each test session.

A licensed phlebotomist inserted a saline-lock intravenous line into the dominant or non-dominant hand or forearm daily to evaluate hypothalamic-pituitary-gonadal axis hormones, including serum levels of gonadal hormones (17*β*-estradiol, progesterone and testosterone) and the pituitary gonadotropins luteinizing hormone (LH) and follicle stimulating hormone (FSH). One 10cc mL blood sample was collected in a vacutainer SST (BD Diagnostic Systems) each session. The sample clotted at room temperature for 45 min. until centrifugation (2,000 g for 10 minutes) and was then aliquoted into three 1 ml microtubes. Serum samples were stored at −20 C until assayed. Serum concentrations were determined via liquid chromatography-mass spectrometry (for all steroid hormones) and immunoassay (for all gonadotropins) at the Brigham and Women’s Hospital Research Assay Core.

### fMRI data acquisition and preprocessing

The participant underwent a daily magnetic resonance imaging scan on a Siemens 3T Prisma scanner equipped with a 64-channel phased-array head coil. First, high-resolution anatomical scans were acquired using a T1-weighted magnetization prepared rapid gradient echo (MPRAGE) sequence (TR = 2500 ms, TE = 2.31 ms, TI = 934 ms, flip angle = 7°; 0.8 mm thickness) followed by a gradient echo fieldmap (TR = 758 ms, TE1 = 4.92 ms, TE2 = 7.38 ms, flip angle = 60°). Next, the participant completed a 10-minute resting-state fMRI scan using a T2 -weighted multiband echoplanar imaging (EPI) sequence sensitive 468 to the blood oxygenation level–dependent (BOLD) contrast (TR = 720 ms, TE = 37 ms, flip angle = 56°, multiband factor = 8; 72 oblique slices, voxel size = 2 mm). In an effort to minimize motion, the head was secured with a custom, 3D-printed foam head case (https://caseforge.co/) (days 8–30 of Study 1). Overall motion (mean frame-wise displacement) was negligible, with fewer than 130 microns of motion on average each day.

Initial preprocessing was performed using the Statistical Parametric Mapping 12 software (SPM12, Wellcome Trust Centre for Neuroimaging, London) in MATLAB. Functional data were realigned and unwarped to correct for head motion and the mean motion-corrected image was coregistered to the high-resolution anatomical image. All scans were then registered to a subject-specific anatomical template created using Advanced Normalization Tools (ANTs) multivariate template construction. A 5 mm full-width at half-maximum (FWHM) isotropic Gaussian kernel was subsequently applied to smooth the functional data. Further preparation for resting-state functional connectivity was implemented using in-house MATLAB scripts. Global signal scaling (median = 1,000) was applied to account for fluctuations in signal intensity across space and time, and voxelwise timeseries were linearly detrended. Residual BOLD signal from each voxel was extracted after removing the effects of head motion and five physiological noise components (CSF + white matter signal). Motion was modeled using a Volterra expansion of translational/rotational motion parameters, accounting for autoregressive and nonlinear effects of head motion on the BOLD signal. All nuisance regressors were detrended to match the BOLD timeseries. Functional network nodes were defined based on a 400-region cortical parcellation and 15 regions from the Harvard–Oxford subcortical atlas. For each day, a summary timecourse was extracted per node by taking the first eigenvariate across functional volumes. These regional timeseries were then decomposed into several frequency bands using a maximal overlap discrete wavelet transform. Low-frequency fluctuations in wavelets 3–6 (0.01–0.17 Hz) were selected for subsequent connectivity analyses. Finally, we estimated the spectral association between regional timeseries using magnitude-squared coherence: this yielded a 415 x 415 functional association matrix each day, whose elements indicated the strength of functional connectivity between all pairs of nodes (FDR-thresholded at q *<* .05).

### Dynamic community detection and anal-ysis

Communities in resting-state connectivity were identified by maximizing multislice modularity, given by

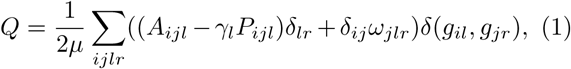

where *µ* is the total edge weight in the network, *i* and *j* index nodes in slices *l* and *r, A* is the adjacency matrix containing edge weights between nodes and slices, *γ* is the structural resolution parameter, *P* is the optimization null model adjacency matrix, δ is the Kronecker delta, *ω* is the temporal resolution parameter, and *g* is the community assignment index [24]. Community assignments that maximize modularity were determined 50 times over a grid of parameter values (*γ,ω*) = [.95, 1.1]x[.8, 1.2] using the genlouvain function from Jeub et al. in MATLAB 2019a [52]. From these community assignments, the consensus partition for each parameter combination was determined using the consensus similarity function from the Networ<u>k</u> Connectivity Toolbox (NCT, http://commdetect.weebly.com/).

Node flexibility is defined as the proportion of times a node changes community assignment out of all possible opportunities to change its assignment. Thus, a flexibility value of 1 indicates that a node changes community membership at every time step and a value of 0 indicates that it never changes communities. Partition significance, node flexibility, and persistence were also calculated using functions from the NCT [36]. Cross-covariance values were calculated and statistical tests were performed using built-in MATLAB functions.

## Acknowledgments

This work was supported by the Brain and Behavior Research Foundation, the California Nanosystems Institute, the Hellman Family Fund, the Rutherford B. Fett Fund, the David and Lucile Packard Foundation and the Institute for Collaborative Biotechnologies through contract no. W911NF-09-D-0001 from the U.S. Army Research Office. Thanks to Mario Mendoza for phlebotomy and MRI assistance. We would also like to thank Shuying Yu, Courtney Kenyon, Maggie Hayes, and Morgan Fitzgerald for assistance with data collection.

