## Supplemental Figures for "Dynamic community detection reveals transient reorganization of functional brain networks across a female menstrual cycle"

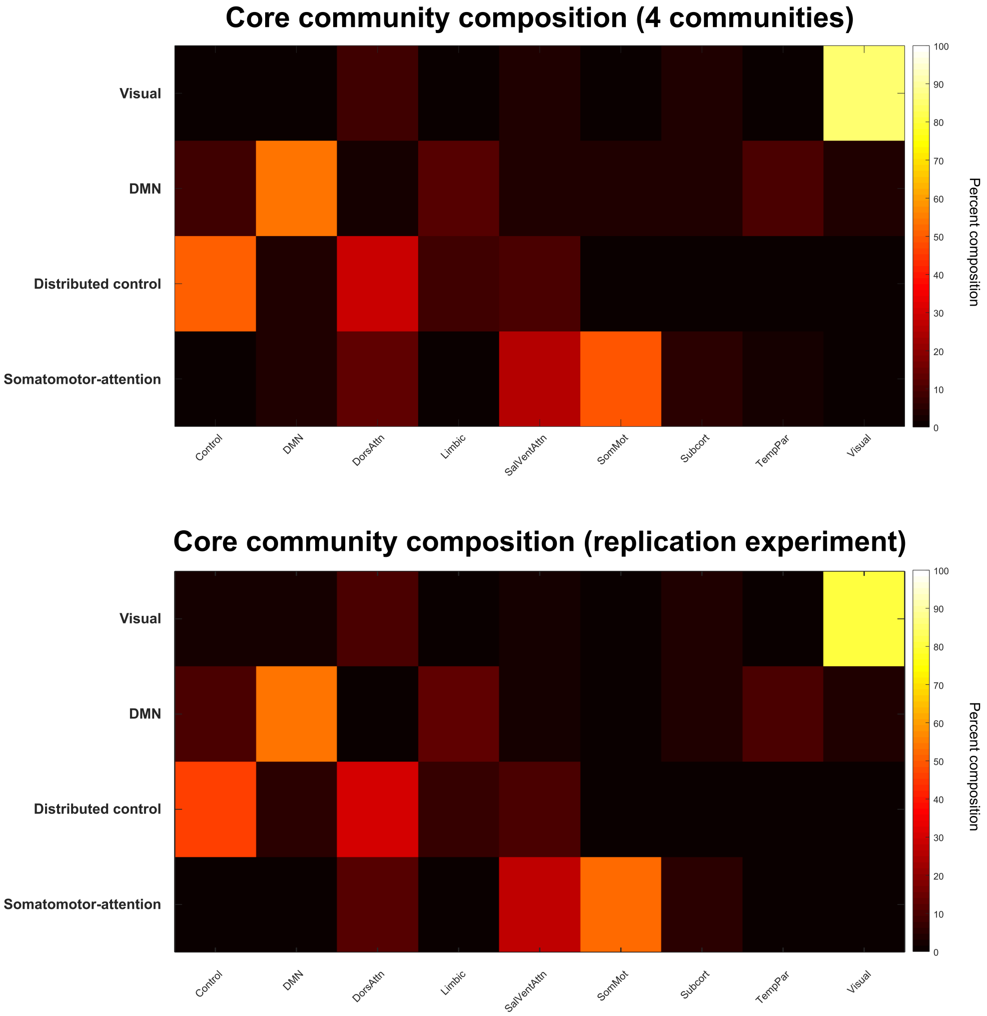

**S1 Fig. Core community compositions are consistent across dynamic community detection parameter choices.** Shown are heat maps of the core community compositions for the original experiment (top) and hormone suppression replication experiment (bottom). In each case, community compositions are averaged over the set of parameter combinations which result in four communities being identified ($.8 < \omega < 2$, $.995 < \gamma < 1.01$). The compositions shown here are nearly identical to those shown in Fig. 3A, which is the core community composition identified at the standard parameter combination for the dynamic community detection algorithm ($\omega = \gamma = 1$).

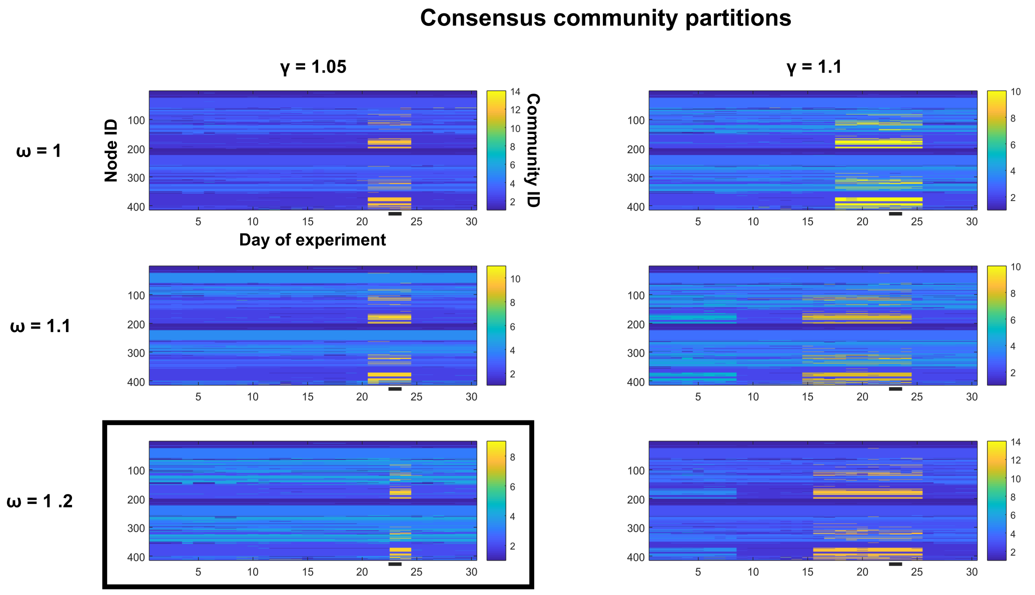

**S2 Fig. Functional communities exhibit transient reorganization around the ovulatory window over a range of dynamic community detection parameter values.** Shown are the consensus community partitions for a range of dynamic community detection algorithm parameters ($1 < \omega < 1.2$, $1.05 < \gamma < 1.1$). Y-axis values indicate node identity, x-axis values indicate the day of experiment, and color indicates community membership. Ovulation occurs on day 23 (black underline). The partition outlined in black (bottom left) is the basis of the quantitative analysis in the main text and is shown in Fig. 4B. In each case displayed here, a new subcommunity with a consistent composition splits from the default mode core during the ovulatory window but rejoins after day 25 at the latest, concurrent with a sharp decline in estradiol, LH, and FSH (Fig. 6B).

| Node Label | Network | Subnetwork | xMNI | yMNI | zMNI |
|---|---|---|---|---|---|
| 17Networks_LH_DefaultB_Temp_1 | DMN | DMN B | -44 | 12 | -34 |
| 17Networks_LH_DefaultB_Temp_2 | DMN | DMN B | -54 | -2 | -30 |
| 17Networks_LH_DefaultB_Temp_3 | DMN | DMN B | -62 | -18 | -20 |
| 17Networks_LH_DefaultB_Temp_4 | DMN | DMN B | -56 | -8 | -14 |
| 17Networks_LH_DefaultB_Temp_5 | DMN | DMN B | -60 | -34 | -4 |
| 17Networks_LH_DefaultB_Temp_6 | DMN | DMN B | -52 | -22 | -6 |
| 17Networks_LH_DefaultB_IPL_1 | DMN | DMN B | -46 | -58 | 20 |
| 17Networks_LH_DefaultB_IPL_2 | DMN | DMN B | -56 | -54 | 30 |
| 17Networks_LH_DefaultB_PFCd_1 | DMN | DMN B | -4 | 52 | 28 |
| 17Networks_LH_DefaultB_PFCd_2 | DMN | DMN B | -14 | 58 | 30 |
| 17Networks_LH_DefaultB_PFCd_3 | DMN | DMN B | -22 | 50 | 32 |
| 17Networks_LH_DefaultB_PFCd_4 | DMN | DMN B | -8 | 42 | 52 |
| 17Networks_LH_DefaultB_PFCd_5 | DMN | DMN B | -12 | 24 | 60 |
| 17Networks_LH_DefaultB_PFCd_6 | DMN | DMN B | -6 | 10 | 64 |
| 17Networks_LH_DefaultB_PFCl_1 | DMN | DMN B | -40 | 20 | 48 |
| 17Networks_LH_DefaultB_PFCl_2 | DMN | DMN B | -42 | 8 | 48 |
| 17Networks_LH_DefaultB_PFCv_1 | DMN | DMN B | -36 | 22 | -16 |
| 17Networks_LH_DefaultB_PFCv_2 | DMN | DMN B | -36 | 36 | -12 |
| 17Networks_LH_DefaultB_PFCv_3 | DMN | DMN B | -46 | 32 | -10 |
| 17Networks_LH_DefaultB_PFCv_4 | DMN | DMN B | -48 | 28 | 0 |
| 17Networks_LH_DefaultB_PFCv_5 | DMN | DMN B | -54 | 20 | 12 |
| 17Networks_RH_DefaultB_Temp_1 | DMN | DMN B | 64 | -24 | -8 |
| 17Networks_RH_DefaultB_Temp_2 | DMN | DMN B | 64 | -38 | 0 |
| 17Networks_RH_DefaultB_AntTemp_1 | DMN | DMN B | 50 | 8 | -32 |
| 17Networks_RH_DefaultB_PFCd_1 | DMN | DMN B | 6 | 58 | 30 |
| 17Networks_RH_DefaultB_PFCd_2 | DMN | DMN B | 16 | 52 | 36 |
| 17Networks_RH_DefaultB_PFCd_3 | DMN | DMN B | 4 | 44 | 40 |
| 17Networks_RH_DefaultB_PFCd_4 | DMN | DMN B | 14 | 38 | 52 |
| 17Networks_RH_DefaultB_PFCd_5 | DMN | DMN B | 12 | 20 | 62 |
| 17Networks_RH_DefaultB_PFCv_1 | DMN | DMN B | 34 | 22 | -18 |
| 17Networks_RH_DefaultB_PFCv_2 | DMN | DMN B | 48 | 32 | -8 |
| 17Networks_RH_DefaultB_PFCv_3 | DMN | DMN B | 54 | 24 | 6 |
| 17Networks_LH_ContC_pCun_1 | Control | Control C | -10 | -70 | 32 |
| 17Networks_LH_ContC_pCun_2 | Control | Control C | -10 | -78 | 46 |
| 17Networks_LH_ContC_pCun_3 | Control | Control C | -4 | -64 | 52 |
| 17Networks_LH_ContC_Cingp_1 | Control | Control C | -6 | -40 | 24 |
| 17Networks_LH_ContC_Cingp_2 | Control | Control C | -4 | -22 | 30 |
| 17Networks_RH_ContC_pCun_1 | Control | Control C | 16 | -64 | 28 |
| 17Networks_RH_ContC_pCun_2 | Control | Control C | 14 | -72 | 40 |
| 17Networks_RH_ContC_pCun_3 | Control | Control C | 6 | -64 | 44 |
| 17Networks_RH_ContC_pCun_4 | Control | Control C | 8 | -50 | 44 |
| 17Networks_RH_ContC_pCun_5 | Control | Control C | 8 | -72 | 52 |
| 17Networks_RH_ContC_Cingp_1 | Control | Control C | 8 | -44 | 20 |
| 17Networks_RH_ContC_Cingp_2 | Control | Control C | 6 | -28 | 28 |

**S3 Fig. Node identities from the Schaefer functional-anatomical atlas.** Shown here are the node identities for regions in the Control C and DMN B subnetworks. Control C nodes are identified as being highly flexible over the entire course of the menstrual cycle. Within-network connectivity between nodes in DMN B increases around the ovulatory window, resulting in a transient bifurcation of the default mode core community (Fig. 5).

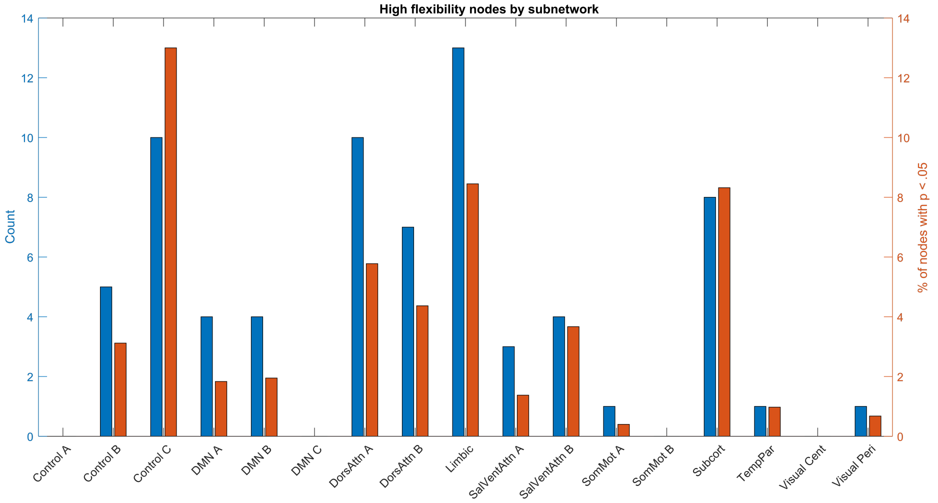

**S4 Fig. Functional-anatomic subnetworks have distinct flexibility profiles.** Within functional-anatomical networks, nodes belonging to different subnetworks exhibit different flexibility trends. Specifically, Control C subnetwork nodes are the most likely to be highly flexible within the Control network, suggesting a specific "integrator" role for this subnetwork.

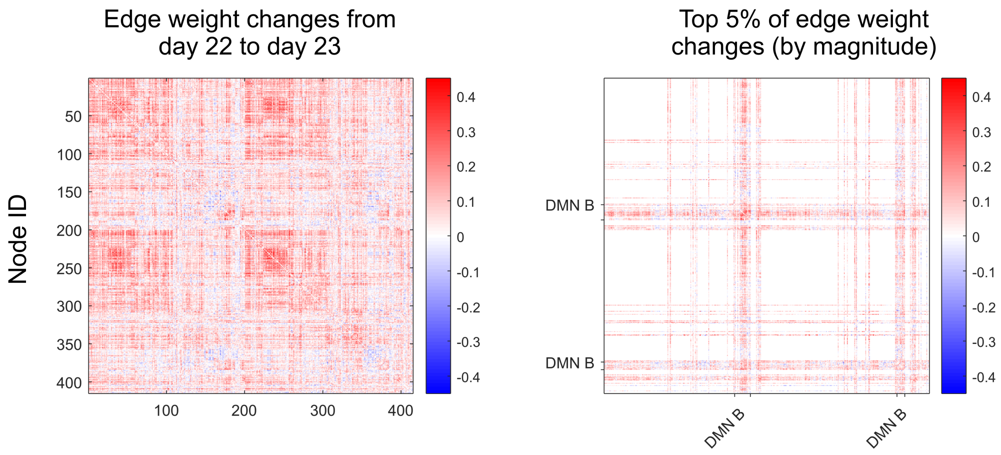

**S5 Fig. Strong changes in functional connectivity during the ovulatory window are localized to a default mode subnetwork.** Shown here are the differences in edge weights (magnitude-squared coherence values) between nodes on days 22 and 23, with color indicating value. Positive values indicate an increase in edge weight from day 22 to 23. On the right, only the top 5% of changes by magnitude are displayed. As demonstrated by the extent of edge weights present in the rows and columns labeled DMN B, it can be seen that within-subnetwork changes in edge strength are responsible for the observed community reorganization during the ovulatory window.
